# *Centella asiatica* improves sleep quality and quantity in aged mice

**DOI:** 10.1101/2025.02.01.636070

**Authors:** Laura Dovek, Carolyn E. Tinsley, Katelyn Gutowsky, Kayla McDaniel, Zoe Potter, Matthias Ruffins, Noah E.P. Milman, Claire Wong, Amala Soumyanath, Nora E. Gray, Miranda M. Lim

**Author notes:** denotes equal contribution (Laura Dovek, Carolyn E. Tinsley). Correspondence should be addressed to: Miranda Lim Portland VA Health Care System Mailcode: P3-RD42 3710 SW US Veterans Hospital Rd Portland, OR 97239.

## Abstract

Age-related sleep disruption is common in older adults. Not only does the total amount of time spent in sleep decline, but the number of arousals during sleep increases with age. As sleep is important for both memory consolidation and to prevent neurodegenerative pathology, this decline in sleep and/or sleep consolidation may underlie age-related cognitive decline and dementias. Furthermore, treatment of sleep disruption can improve quality of life. However, few interventions have successfully reversed age-related sleep decline. Extracts from the plant *Centella asiatica* have demonstrated neuroprotective effects in human, rodent, and fly models of aging and neurodegenerative diseases, and is a promising intervention for dementias, yet little is known about how these extracts affect sleep patterns. Here, we administered *Centella asiatica water extract* (*CAW)* dosed or control chow to male and female C57BL6/J mice aged 18 months. Effects on sleep composition were determined using electrodes that recorded EEG and EMG signals. We found that CAW dosed chow (1000 mg/kg/day) increased REM sleep time in aged male mice and decreased the number of arousals during sleep observed in aged females, compared to age- and sex-matched controls. We conclude that CAW administered in food has a moderate, sex-dependent effect on sleep quantity and quality.

**Statement of Significance:** Sleep declines with age and may underline age-related cognitive changes. However, few interventions have successfully reversed age-related sleep and cognitive decline. This study found that botanical extract from the plant *Centella asiatica* increased total REM sleep time in aged male mice, and decreased sleep fragmentation in aged female mice, compared to age- and sex-matched controls. Whether these moderate, sex-dependent effect sizes on sleep in aged mice are impactful enough to affect cognition, quality of life, and/or neurodegenerative pathology could be explored in future studies.

## Introduction

Sleep plays a crucial role in maintaining both physical and mental well-being. Sleep acts as much needed time for physiologic restorative processes, memory consolidation and regulation of several bodily functions ^1–4^. With age, sleep undergoes significant changes including alterations in both the quality and quantity of sleep. Older adults often have increased difficulty getting into deep sleep and impaired sleep maintenance resulting in less total sleep time compared to younger individuals ^5^. Poor sleep is connected with many adverse health outcomes relevant to the aging population including, impairments in memory and cognition. Sleep is necessary for the processing of newly acquired information and facilitating consolidation of memories into long-term storage ^6–9^. In rodents, both the quality and quantity of sleep is important for learning and memory with artificial enhancement or deprivation of sleep improving or impairing performance, respectively, on a variety of memory tasks ^2,6,10–12^.

Additionally, with aging, comes an increased likelihood of the development of neurodegenerative and psychiatric disorders including Alzheimer’s and Parkinson’s Disease ^13–15^. This relationship is complex and multidirectional; aging and neurodegenerative conditions can result in sleep difficulties as well as memory impairment, while disrupted sleep can negatively impact memory and precede or even exacerbate the development of neurodegenerative diseases. While it is unclear whether sleep disturbances are the cause or simply a symptom of underlying disease; sleep, aging, and neurodegenerative disorders are highly intertwined Given the importance of sleep quality on several aspects of our health, it is necessary to identify strategies to ameliorate subjective sleep symptoms which are troublesome in older adults and to enhance sleep to decelerate progression of neurodegenerative disorders. *Centella asiatica* is a botanical whose bioactive compound extracts are shown to target several signaling pathways which are associated with inflammation and aging ^16–18^. *Centella asiatica* is actively used for numerous health purposes including neuroprotection, dermatological protection, improving mental health, organ protection, and has shown protective effects following traumatic brain injuries as well as benefits in Alzheimer’s and Parkinson’s disease animal models ^19–25^. *Centella asiatica* has the potential to be a neuroprotectant therapy as its targets are believed to regulate anti-aging activity by modulating the previously mentioned signaling pathways resulting in decrease in inflammation and direct elimination of reactive oxygen species (ROS) ^18,26^. In humans, *Centella asiatica* has been shown to improve mental state exam scores in elderly individuals with mild cognitive impairment as well as other common age-related conditions such as improvement in mood disorders, digestion, and insomnia ^22^. Furthermore, *Centella asiatica* treatment has been shown to improve memory and promote antioxidative signaling in mice models ^20^. How *Centella asiatica* influences sleep architecture, and if this could explain the cognitive improvements observed following *Centella asiatica* treatment, has yet to be determined.

The present study evaluated water extracted Centella asiatica’s (CAW) influence on quality and quantity of sleep. Using younger (3m) and older (18m) C57Bl/6 mice, we first establish a baseline of aging’s effects on both female and male sleep patterns using control animals. Then, we assess a CAW diet’s influence on sleep patterns of aged mice. Finally, we utilize a novel object recognition task to see if CAW can restore deficits in recognition memory following sleep disruption.

## Methods

### Experimental Design

In this study, we explored the effects of water extracted *Centella asiatica (CAW)* on sleep in aged, 18-months old (18m) C57BL/6 mice by analyzing sleep patterns through EEG/EMG recordings. Mice were assigned a treatment or control group before experimentation and were grouped as follows: n=8 control/18m/female, n=9 control/18m/male, n=8 CAW/18m/female, and n=9 CAW/18m/male. Additional cohorts of younger, 3-month old (3m) mice were given control chow (n= 8 control/3m/female and n= 9 control/3m/male) in order to describe the effects of age on sleep in this population.

On day 1, the treatment group was given unrestricted access to CAW-dosed chow (animals consumed on average 1000 mg/kg/day), and water. Our control group was also given unrestricted access to control chow and water. On day 16, mice underwent stereotaxic implantation of indwelling electrodes. On day 20, sleep was recorded for 3 days while they continued to have unrestricted access to CAW-dosed chow or control chow. On day 24, mice were untethered and humanely euthanized using 5% inhaled isoflurane followed by cervical dislocation.

### Subjects

Male and female C57BL/6J mice were acquired from the NIA Aged Rodent Colony. Mice were housed in same-sex pairs until EEG/EMG electrode implantation and then single-housed for the remainder of the study. Males and females were always run in separate experimental cohorts, with no overlap in housing or procedures. As such, sex was not used as a between group factor in analysis. Mice were kept in a climate-controlled environment while exposed to a 12-hour light-dark cycle (lights on a 0700 h). Our treatment and control groups were provided *ad libitum* access to CAW-dosed chow or control chow for 24 days, respectively. Water was provided *ad libitum.* All experiments and procedures were reviewed and approved by the Institutional Animal Care and Use Committee (IACUC) at VA Portland Health Care System.

### CAW Dosed Chow

CAW was prepared as previously described by our group^27^. Briefly, 4kg of *Centella asiatica* (dried aerial parts) purchased through Oregon’s Wild Harvest (Redmond, OR, USA) was boiled for 90 minutes in 50L of deionized water. The cooled mixture was filtered to remove plant debris and the filtrate was frozen in aluminum baking trays and lyophilized in 3 batches (total weight of 820 g) all of which were used during the course of the present study. Voucher samples of the starting plant material are stored at the Oregon State University Herbarium (Voucher number OSC-V265416) and in our laboratory at Oregon Health & Science University.

Diets containing CAW were prepared by Dyets Inc. (Bethlehem PA) by mixing CAW at 1.0 % w/w with the AIN-93M diet components until a homogenous distribution was achieved. This concentration of CAW was estimated to deliver an approximate exposure of 1000mg/kg/d to the mice. Inclusion of CAW in the rodent diets at the required levels was verified using liquid chromatography multiple reaction monitoring mass spectrometry (LC-MRM-MS) and previously reported^28^. Control AIN-93M diet without CAW was also acquired from Dyets. Both diets was sterilized by gamma irradiation (5.0-20.0 kGy; Sterigenics, Oak Brook, IL, USA). Chow was stored at -4 C until used.

### Surgical Implantation of EEG/EMG Electrode

EEG and EMG electrodes were custom built as described previously^29,30^. Female pin headers (853-93-100-10-001000, DigiKey) were coated with liquid solder^60^ flux (10-4202, GC Electronics) before attaching seven, 3-5mm silver wire (782500, A-M systems) with silver solder (SN60PB40, Kester) ^30^. Stainless steel screws (Precision Screws and Parts) were coated with stainless steel soldering flux liquid (23904, LA-CO) and attached to the top four silver wires with stainless steel solder (SN96.5AG03.5, Kester). The electrode was then de-fluxed using flux-off (ESZ835B, Chemtronics). The electrodes were tested with a voltmeter before being coated with Corona Dope (10-4702, GC Electronics) and before surgical implantation.

Anesthesia was induced at 5% isoflurane/1L/m oxygen and maintained between 1-3% isoflurane/1L/m oxygen. Mice were transferred to a stereotaxic apparatus and then secured to a bite bar and stabilizing ear bars. The eyes were protected with Puralube ointment (NDC 24208-780-55, Bausch + Lomb). The hair was removed by using surgical scissors and the scalp was disinfected with iodine, alcohol and swabbed with 2.5% lidocaine ointment. Once the skin was dry, a 1.5cm incision was made down the midline skin tissue. Sterile swabs with 0.3% hydrogen peroxide were used on the skull to remove the periosteum and reveal bregma and lambda markings. Four holes were drilled into each brain quadrant by referencing bregma and lambda markings. Two silver wires acting as the EMG wires were threaded through the neck muscle before mounting the electrode box to the skull with super glue. The four silver screws were implanted in each burr hole by manually rotating ¼ of a turn. When the screws were secured, the electrode box, EEG/EMG wires, and EEG screws were then coated in a 1:1 ratio of dental cement and methyl methacrylate (M55909, Sigma-Aldrich). Sutures were placed in the skin above the neck muscle to ensure proper healing. Antibiotic ointment was coated over the surgical site and around the electrode box. Post-operative analgesia was achieved by administering cherry flavored children’s acetaminophen in drinking water as needed.

### Sleep Experiments

EEG and EMG signals were acquired using the Biopac MP160 Data Acquisition System (Biopac Systems, Inc., Goleta, CA). Mice were hooked up to the tethered recording system in their home cages 4 days after surgery, recorded for 72 hours and monitored daily. Mice received Hydrogel, CAW-dosed chow for the treatment group, and control chow for the control group for the entire duration of sleep recording. EEG/EMG recordings were collected as Acknowledge files and converted to KCD files to be analyzed using SleepSign (Kissei Comtec, Nagano, Japan) software. Files were divided into 4 second epochs and each epoch was assigned to one of 3 stages: wake, REM, or NREM. Staging was determined by identifying key characteristics associated with EEG and EMG wave formations with each epoch manually reviewed by trained researchers blind to age, group, and sex. Wake was determined by observing low-voltage and fast EEG waveforms with high-amplitude EMG wave patterns. In NREM sleep, EEG waveforms have a higher amplitude and EMG signals have lower amplitude. EEG signals in REM sleep mimic those in wake but the differentiating factor is that the EMG signal amplitudes are the lowest. Three mice (n=1 CAW/18m/F, n=2 Control/18m/F, n=1 CAW/18m/M were excluded from this experiment due to surgical complications or poor signal quality.

### Sleep Deprived Novel Object Recognition Task

In order to test the effects of CAW on learning and memory, a separate cohort of mice receiving 1% CAW in the chow underwent sleep deprived novel object recognition testing (SD-NORT) as described previously ^31^. Briefly, in an arena 38 x 38 x 64 cm high constructed of white acrylonitrile butadiene styrene, after two 10-minute habituation sessions, animals are exposed to two identical objects for 10-minute training sessions, once an hour over a 3 hour period. After the final training session, mice were kept awake for 6 consecutive hours by intermittent brushing with a small paintbrush. The testing phase occurred 24 hours after the final training session where one of the objects was replaced with a novel object and the time spent exploring the familiar and novel objects over 5 minutes was evaluated via a camera placed above the arena, interfaced with ANYmaze video tracking system (Stoelting Co, Wood Dale, IL). Objects were all approximately 4-6 cm tall and included small plastic boxes, blocks, cups and saltshakers. The use of objects was counterbalanced so that each object was used equally as either the familiar or the novel object. Videos were manually scored by an investigator blinded to the treatment conditions to assess time the mice were exploring each object without climbing on it.

### Statistical Analysis

Conditions were met for parametric testing. Data was analyzed using GraphPad Prism 10.2.3. Sleep metrics for the middle 24 hours of recording for each animal were summed for a 12 hour light and 12 hour dark period. Unpaired t-test, 2-Way and 3-Way ANOVAs, when appropriate, were conducted to compare the effects of age, light/ dark portions of the light cycle and/or treatment. For visualization purposes, data was aggregated into 2-hour time bins to appreciate the effects throughout the course of the day. Significant or trending main effects and interactions were followed up using post-hoc Šídák’s multiple comparisons test (two tailed). An alpha value of 0.05 was used to determine significance.

## Results

### Sleep and Aging

#### Sleep Duration

In order to examine the effects of aging on sleep, we first compared sleep in animals fed control chow that were either 3 months or 18 months of age. There was no significant effect of age on total sleep time (NREM + REM) in males or females (between subjects effect of age: males: F (1, 26) = 0.002, p=0.297; females: F (1, 26) = 1.43 p=0.242). As expected, there was a significant effect of light cycle with both males and females exhibiting less sleep, or more wakefulness during the dark portion of the light cycle (within subjects effect of light: males: F (1, 26) = 26.06, p<0.0001; females: F (1, 26) = 112.20, p<0.0001) (Fig 1A). Age did not influence total time spent in NREM in males nor females (between subjects effect of age: males: F (1, 26) = 0.10, p=0.75; females: F (1, 26) = 0.82, p=0.373) while light did influence the amount of time both sexes spent in NREM (within subjects effect of light: males: F (1, 26) = 22.97, p<0.0001; females: F (1, 26) = 94.61, p<0.0001), with both sexes spending more time in NREM sleep during the light portion of the light cycle (Fig 1B).

**Figure 1:**
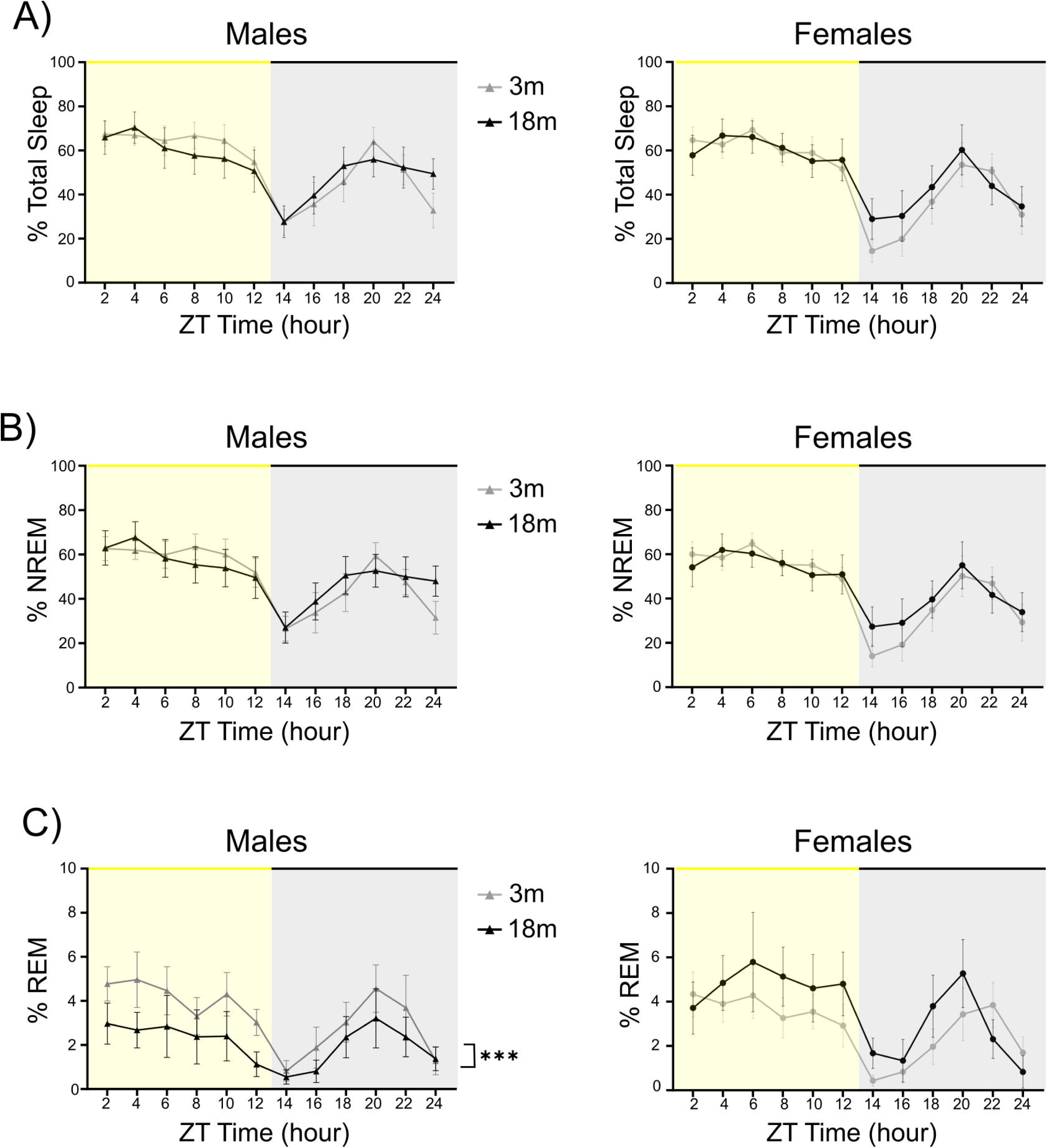
Males show a decrease in REM sleep duration with aging: While age did not influence total sleep time (A) or NREM sleep time (B) in either sex, males spent less time in REM sleep and females trended towards more time in REM sleep at 18 months compared to 3 month old mice of the same sex (C). Data presented as mean ± SEM.

However, we observed sex specific effects of age on REM sleep time. In males, we observed a decrease in time spent in REM with age (between subject effect of age: males: F(1,26) =5.56, p=0.026) (Fig 1C). There was no effect of age on REM sleep duration in females (between subjects effect of age: females: F (1, 26) = 1.58 p=0.220) (Fig 1C). As expected, both males and females spent more time in REM during the light portion of the light cycle compared to the dark portion (within subjects effect of light: males: F (1, 26) = 4.36, p=0.047, females: F(1,26) = 9.53, p=0.005), when nocturnal animals sleep the most. There was no effect of the interaction between age and light portion in both males and females on total sleep time: (interaction of age and light: males: F (1, 26) = 1.11, p=0.302; females: F (1, 26) = 2.13, p=0.157), NREM sleep time: (interaction of age and light: males: F (1, 26) = 0.83, p=0.371; females: F (1, 26) = 2.39, p=0.134), nor REM sleep time (interaction of age and light: males: F (1, 26) = 0.85, p=0.366; females: F (1, 26) = 0.22, p=0.640). These data suggest the effect of age-related REM sleep duration decline is prominent in males.

#### Sleep Fragmentation

Sleep fragmentation was assessed with two different metrics: arousal index (# of sleep-wake arousals/total sleep time), and average sleep (NREM and REM) bout length. Older females experienced more sleep fragmentation than young females as evidenced by increased arousal index (between subjects effect of age: females: F (1, 26) = 11.83, p=0.002) and decreased NREM bout length (between subjects effect of age: females: F (1, 26) = 15.93, p=0.0005) (Fig 2A-B). Holm-Šídák’s multiple comparison tests revealed a trending or significantly increased arousal index in 18m females compared to 3m old females in both the light (p=0.058) and dark (p=0.016) portions of the light cycle (Fig 2A), respectively. This increase in sleep fragmentation was driven by reduced NREM bout lengths in 18m old females compared to 3m (light: p= 0.029, dark: p=0.005) (Fig 2B). There was no effect of light on arousal index (within subjects effect of light: males: (F (1, 26) = 0.59, p=0.451; females: F (1, 26) = 0.04, p=0.844), NREM bout length (within subjects effect of light: males: F(1,26) = 0.79, p=0.383; females: F (1,26)=0.97, p=0.333) nor REM bout length (within subjects effect of light: males: F (1,26)=0.006, p=0.939; females: F (1,26)=0.003, p=0.957) for both females and males. In males, aging neither affected arousal index (between subjects’ effect of age: males: F (1, 26) = 0.0006, p=0.981) nor bout length of NREM (between subjects’ effect of age: males: F (1, 26) = 1.26, p=0.272). Lastly, REM bout lengths saw no effects of aging in both male and female cohorts (between subjects’ effect of aging: males: F (1, 26) = 0.50, p=0.484; females: F (1, 26) = 0.26, p=0.618;) (Fig 2C). There was no effect of the interaction between age and light portion in both males and females on arousal index: (interaction of age and light: males: F (1, 26) = 0.057, p=0.814; females: F (1, 26) = 0.40, p=0.535), NREM bout length: (interaction of age and light: males: F (1, 26) = 0.33, p=0.570; females: F (1, 26) = 0.53, p=0.474), nor REM bout length : (interaction of age and light: males: F (1, 26) = 0.040, p=0.844; females: F (1, 26) = 0.35, p=0.558). All other comparisons were not significant or trending (p>0.1). These data suggest that NREM sleep quality is impaired in aged female mice.

**Figure 2:**
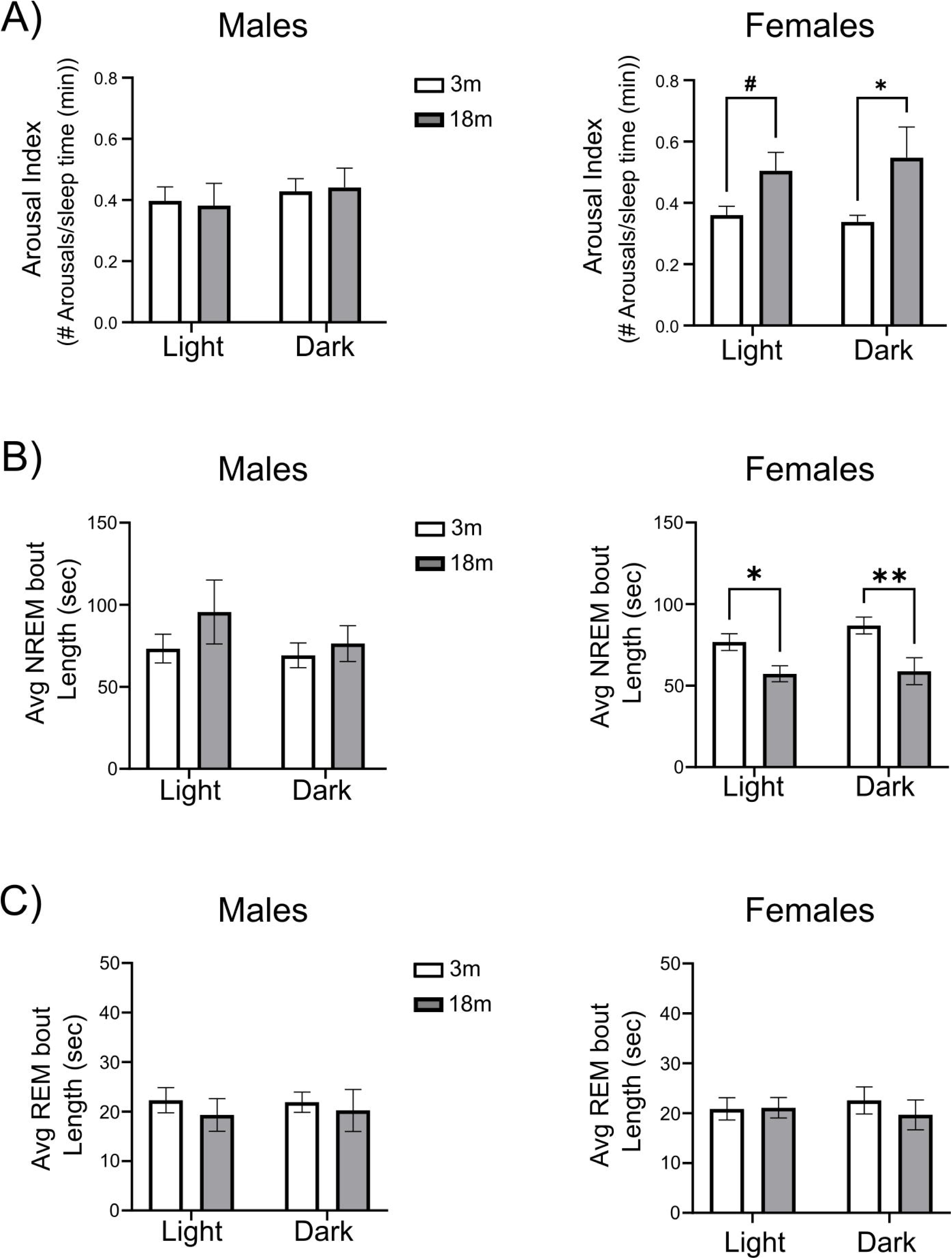
Females experience more fragmented NREM sleep with aging: 18 month old females, but not males, saw an increase in arousal index (#arousals/total sleep time; higher values indicate increased sleep fragmentation) (A) compared to sex matched 3 month old mice. NREM bout lengths were decreased with age in females with no significant effect observed in males (B). There were no changes observed in REM bout length (C). Data presented as mean ±SE, *=p<0.05, **=p<0.01; #=0.058.

#### Sleep and CAW

There was no effect of CAW on total sleep time in males (between subjects effect of CAW: males: F (1,28) = 0.11, p=0.739;) nor females: (between subjects effect of CAW: females: F (1, 22) = 1.47, p=0.239) (Fig 3A-B). All cohorts showed an effect of light with a majority of total sleep taking place during the light portion of the light cycle (within subjects effect of light: males: F (1,28) = 22.37, p<0.0001; females: F (1,22) = 21.79, p=0.0001). The interaction between light phase and treatment condition did not have a significant effect in males and females (between subjects interaction effect: males: F (1, 28) = 0.34, p=0.565; females: F (1, 22) = 0.63, p=0.436).

**Figure 3:**
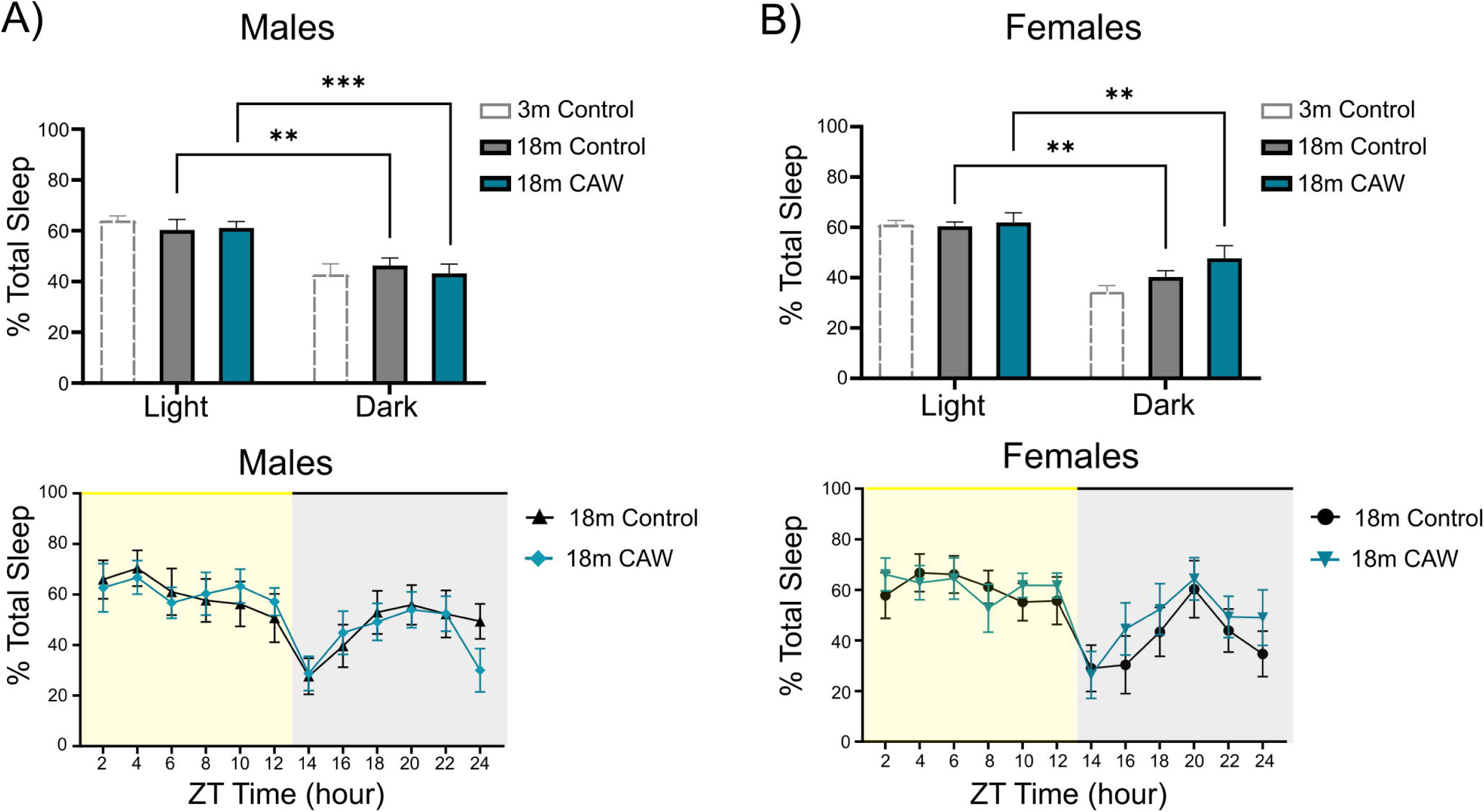
CAW did not influence total sleep time in aged animals: CAW did not influence total sleep time in our 18-month males (A) nor females (B). Data presented as mean ±SE, *=p<0.05, **=p<0.01, ***=p<0.001

There was also no effect of CAW on NREM sleep duration in either sex (between subjects effect of CAW: males: F (1, 28) = 0.77, p=0.387; females: F (1, 22) = 1.50, p=0.234) (Fig 4A-B). Both sexes spent more time in NREM during the light portion of the light cycle (within subjects effect of light: males: F (1,28) = 17.32, p=0.0003; females: F (1,22) = 20.35, p=0.0002). The interaction between light phase and treatment condition did not have a significant effect on NREM sleep in males and females (between subjects interaction effect: males: F (1, 28) = 0.11, p=0.738; females: F (1, 22) = 1.07, p=0.313).

**Figure 4:**
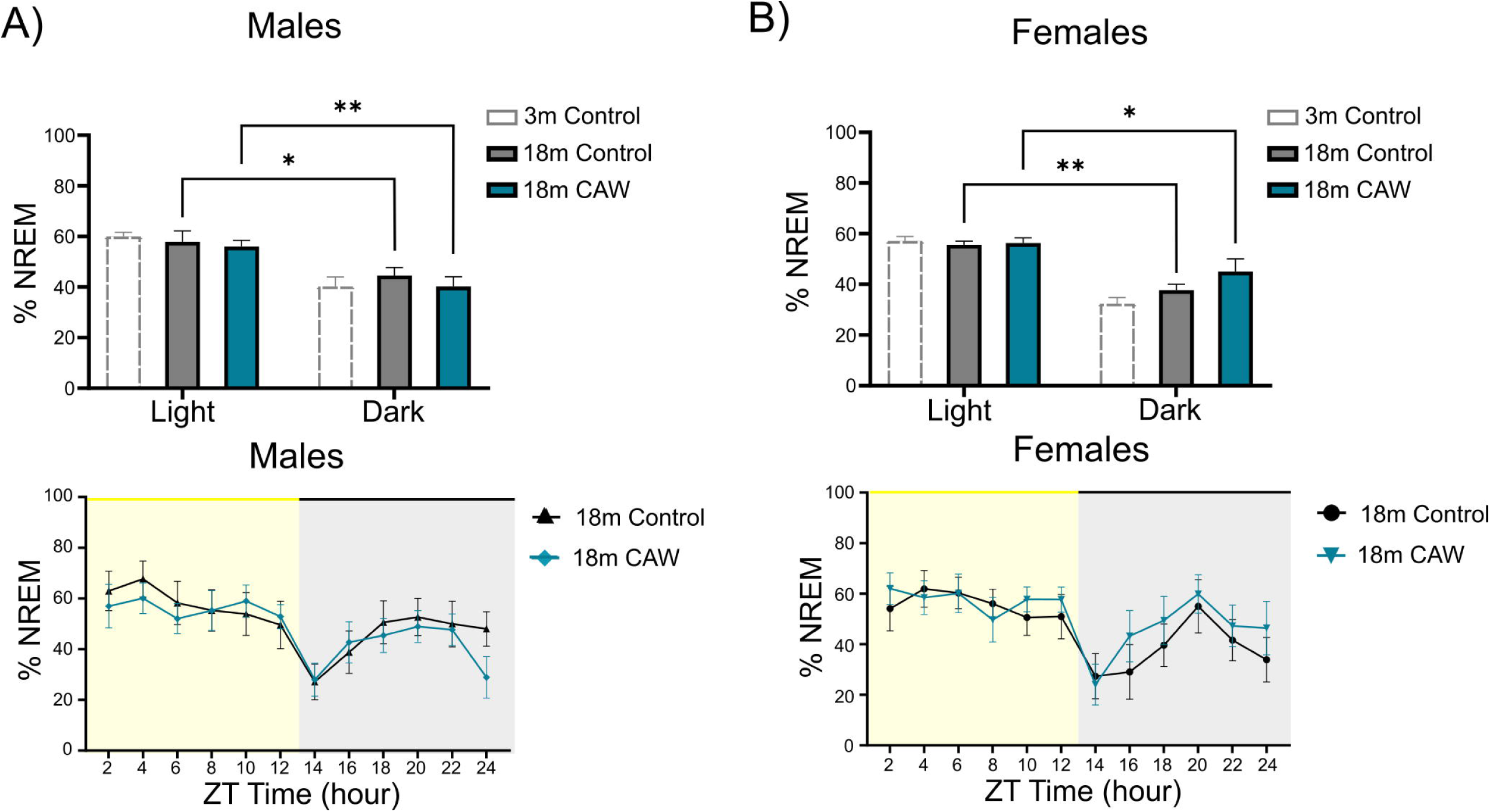
CAW did not affect NREM sleep time: CAW did not influence NREM sleep time in our 18 month males (A) nor females (B). 3 month controls included for visual comparison purposes. Data presented as mean ±SE, *=p<0.05, **=p<0.01

However, CAW resulted in a significant increase in REM sleep time in 18m males (between subjects effect of CAW: males: F (1, 28) = 20.13, p=0.0001) with no significant effect of CAW in females (between subjects effect of CAW: females: F (1, 22) = 0.29, p=0.597) (Fig 5 A,B). For both females and males, there was significantly more time spent in REM sleep during the light portion of the light cycle (within subjects effect of light: males: F (1, 28) = 11.42, p=0.002; females: F (1, 22) = 6.25, p=0.020). Males did show an overall interaction between treatment and light (between subjects interaction effect: males: F (1, 28) = 4.49, p=0.043) where CAW showed the largest increase in REM sleep compared to controls during the light portion of the light cycle when mice are typically asleep(p<0.0001). Females did not show a significant interaction between treatment and light (between subjects interaction effect: females F (1, 22) = 0.481, p=0.495). All other comparisons and interactions were not significant or trending (p>0.1). This data suggests that CAW increases REM sleep time in older males.

**Figure 5:**
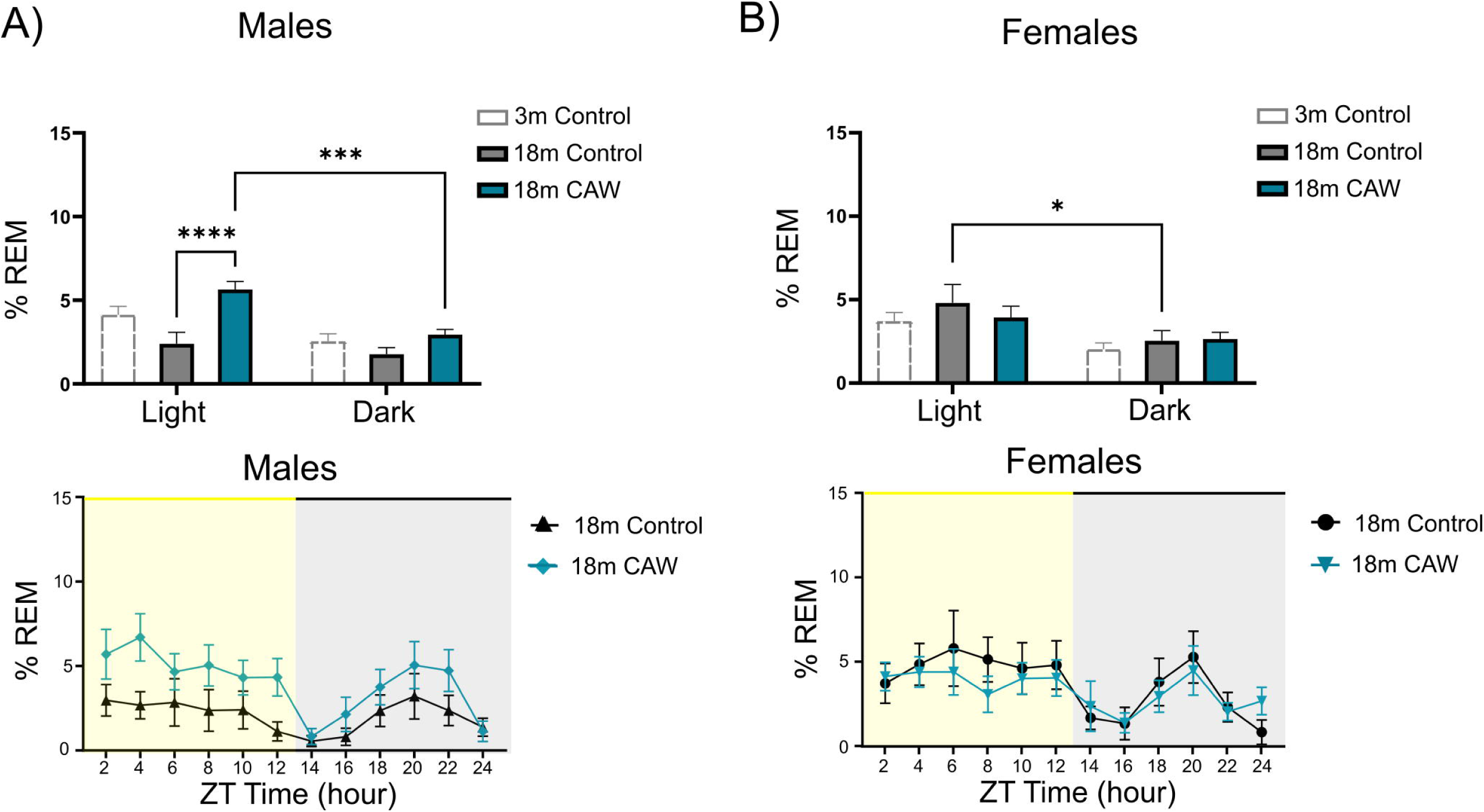
CAW increases REM sleep time in aged males: In our 18m cohort, we saw an increase of REM sleep in males (A) who were fed a CAW diet with no changes observed in 18 month old females (B). 3 month controls included for visual comparison purposes. Data presented as mean ± SE, *=p<0.05, ***=p<0.001, ****=p<0.0001

### CAW’s influence on sleep fragmentation

The addition of CAW significantly reduced the arousal index of aged females (between subjects effect of CAW: females: F (1, 22) = 8.59, p=0.008). Šídák’s multiple comparisons test reveal a significant decrease in arousals between controls and CAW groups during the dark phase (p=0.0336) with the addition of CAW trending to lower arousals between controls and CAW during the light phase (p=0.074). Analysis of arousal index in males revealed no effect of CAW on arousal index (between subjects effect of CAW: males: F (1, 28) = 0.38, p=0.542). Light compared to dark portion of light cycle did not influence arousal index in both males and females (within subjects effect of light: males: F (1, 28) = 0.17, p=0.685; females: F (1, 28) = 0.38, p=0.542) (Fig 6A). There were no interactions between CAW treatment and light phase on arousal index in both male and females (Between subjects interaction effect: males: F (1, 28) = 0.16, p=0.695; females: F (1, 22) = 0.08, p=0.787).

**Figure 6:**
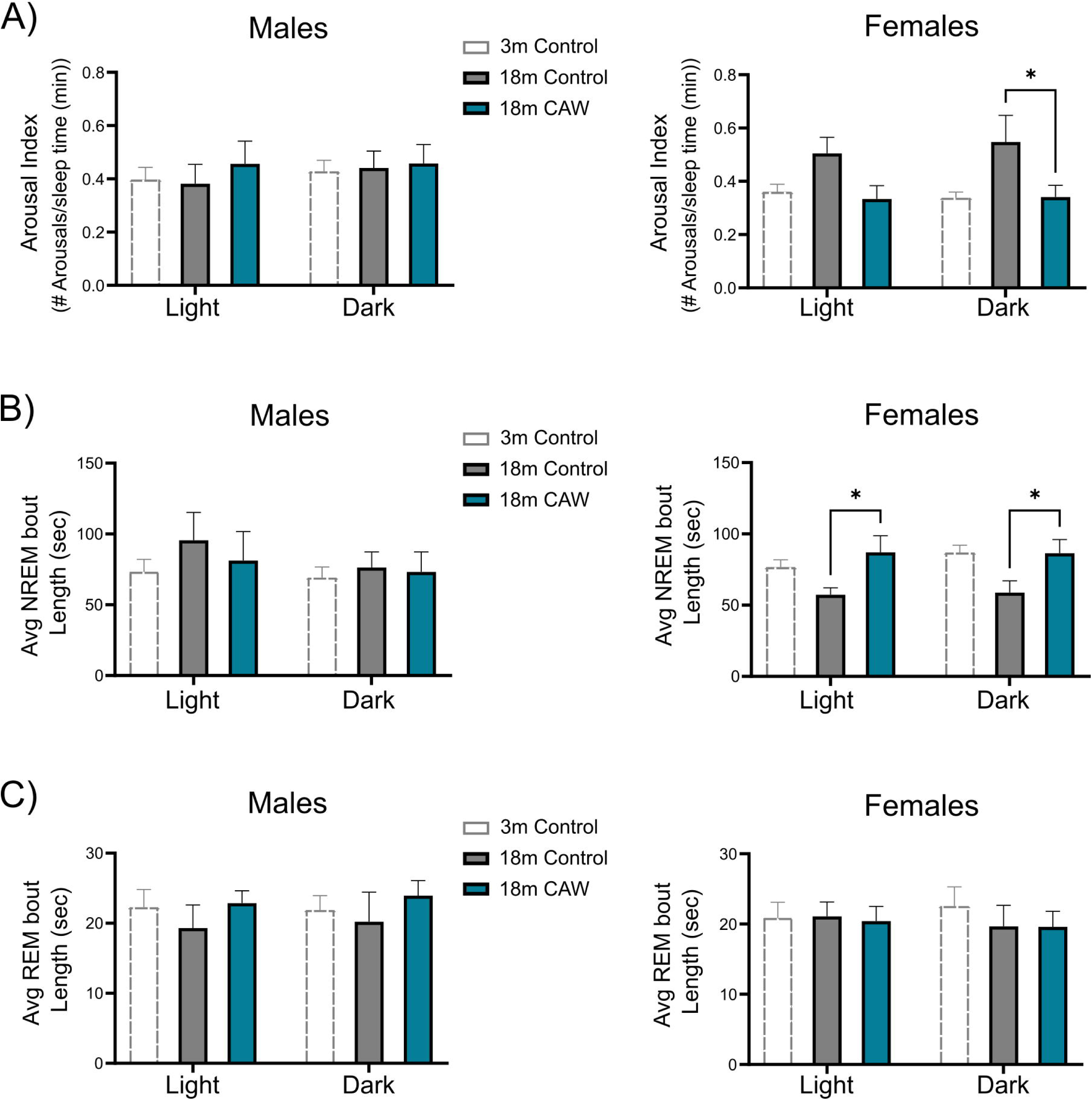
CAW reduces sleep fragmentation in 18m females: CAW reduces arousal index in females but had no influence on males (A). Females but not male NREM bout lengths (B) were increased while REM bout lengths (C) were unaffected as a result of caw in both males and females. Data presented as mean ±SE, *=p<0.05

Lastly, we analyzed the length of NREM and REM bout lengths following the addition of CAW. CAW significantly increased NREM bouts in females (between subjects effect of CAW: females: F (1, 22) = 9.53, p=0.005). Multiple comparisons saw an increase in NREM bout length between controls and CAW groups in the light (p=0.034) and dark (p=0.047) phases (Fig 6B). Males on the other hand saw no changes in NREM bouts as an effect of CAW treatment (between subjects effect of CAW: males: F (1, 28) = 0.28, p=0.603). Furthermore, CAW also did not have an impact on REM bout length in females nor males (between subjects effect of CAW: males: F (1, 28) = 1.45, p=0.238; females: F (1, 22) = 0.02, p=0.877) (Fig 6C). Light cycles did not have an influence on males or females NREM bout lengths (within subjects effect of light: males: F (1, 28) = 0.66, p =0.424; females: F (1, 22) = 0.003, p=0.958) nor REM bout lengths (within subjects effect of light: males: F (1, 28) = 0.11, p=0.745; females: F (1, 22) = 0.23, p=0.640) (Fig 6B-C). There were no interactions observed between light and treatment on NREM bout length (between subjects interaction effect: males: F (1, 28) = 0.11, p=0.738; females: F (1, 22) = 0.01, p=0.911) or REM bout lengths (between subjects interaction effect: males: F (1, 28) = 0. 0009, p=0.977; females: F (1, 22) = 0.02, p=0.900).

### CAW’s influence Cognition following sleep deprivation

Previously, CAW has been shown to improve aged mice performance in several recognition memory and spatial memory tasks. However, it is not known if CAW can still improve performance with an added element of sleep deprivation. To evaluate CAW’s influence on sleep impacted memory, mice underwent behavioral testing. Using the novel object recognition test (NORT), we evaluated recognition memory based on the amount of time an animal spends with a novel object versus a familiar one (Fig 7A). Using a discrimination ratio, or the amount of time spent during the test with the novel object divided by time spent with the familiar object, we failed to observe any differences as a result of aging in both males (p=0.613) and females (p=0.705) (Fig 7B). Furthermore, in our 18m animals, we failed to see any effect of CAW in males (p=0.262) and females (p=0.966). While CAW improves age-related deficits in sleep, we did not observe any age-related impairments in novel object recognition, therefore it is not surprising that CAW did not show improvements.

**Figure 7:**
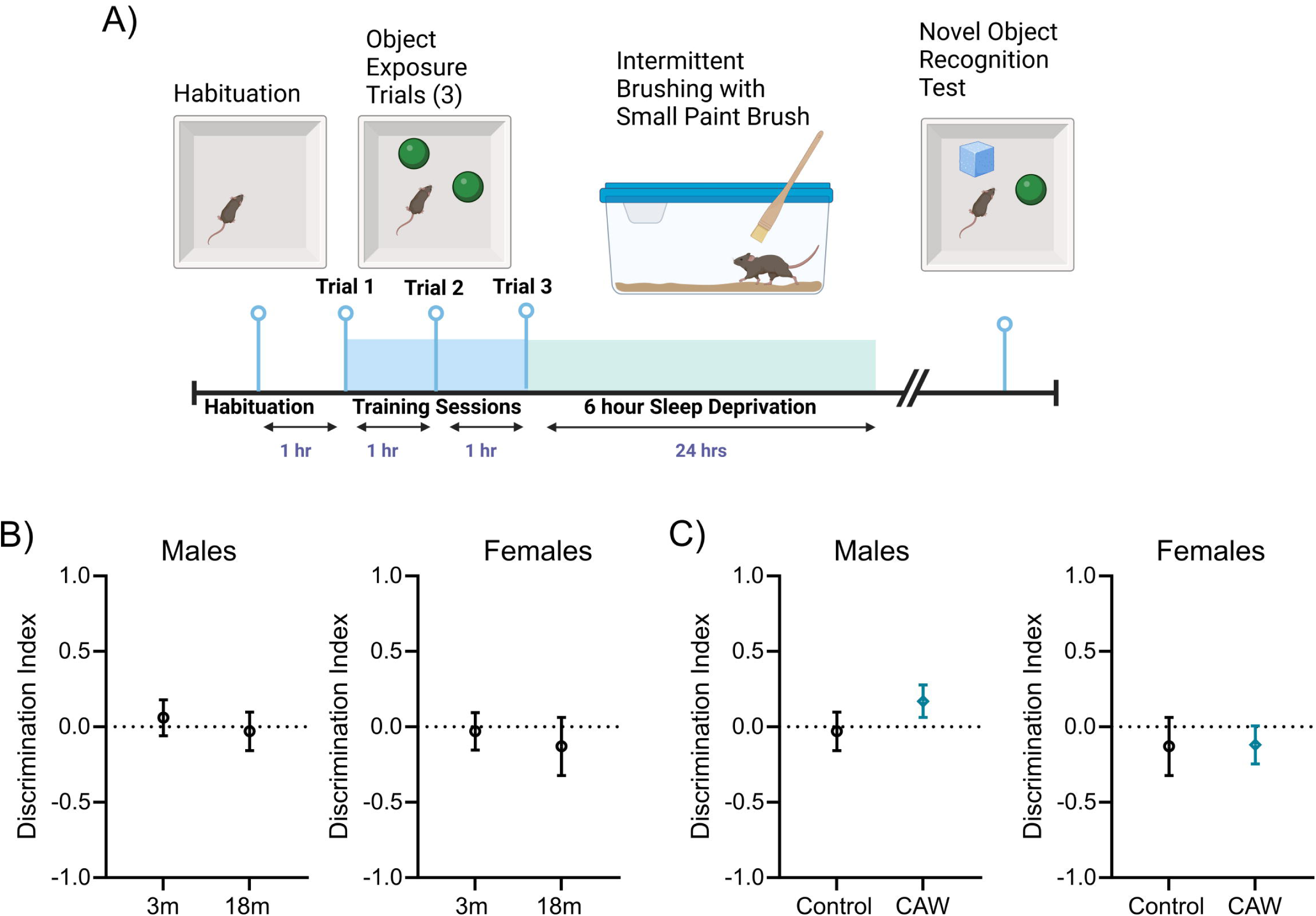
CAW had no effect on performance in Sleep deprived Novel Object Recognition Task: Mice underwent SD NORT training (A) Discrimination Index calculated as (time with novel – time with familiar)/total time with both objects, revealed no difference between young and old (B), and older treated and untreated males and females (C). Figure A created in BioRender. Dovek, L. (2025) https://BioRender.com/i13k750

## Discussion

In these experiments, aged mice were fed control chow or chow dosed with CAW for 20 days followed by EEG/EMG recordings to describe the effects of CAW on sleep. We first describe changes in sleep architecture that occur with age in C57BL/6 mice. We found that, compared to sex-matched 3 month old animals, 18 month old males spend less time in REM sleep (a measure of sleep quantity) and females have shorter average NREM sleep bouts and increased sleep arousals (an indication of sleep quality). We found that 1000mg/kg/day of CAW chow improved these age-related sleep impairments with increased REM sleep time in aged male mice and decreased arousals in aged females compared to age and sex matched mice fed control chow. These findings suggest that CAW improves sleep quantity in older males and sleep quality in older females.

### Aging and its effects on Sleep Architecture

Humans undergo many physiological alterations in normal sleep patterns with aging that are often multifactorial. There appears to be an age-related decrease in the promotion of sleep by the circadian pacemaker as well as a decrease in homeostatic pressure for sleep ^32^. Older adults spend more time in earlier, lighter stages of sleep and less time in deep sleep (SWS and REM)^5,33^. This shift in quality and depth of sleep contributes to decreased sleep maintenance (more frequent awakenings and more fragmented sleep)^32,33^. There is ample evidence of decreased sleep duration and/or quality correlating with decreased mental well-being, impaired cognition, increased fall risk, and increased risk for the development of obesity, diabetes, hypertension, as well as increased likelihood of hospitalizations ^34–43^. Similarly, the inverse, while less studied, is also true. Sleep can be increased using various forms of medication, psychological intervention or simply spending a longer time in bed. Increased sleep and improvements in sleep quality result in improvements in both physical and mental health suggesting a direct beneficial effect of sleep extension ^44,45^.

Mice also show changes in sleep architecture associate with aging. However, results are often mixed and can vary based on genotype and age studied. In the C57Bl/6 mouse, used in this study, most other studies conducted primarily in mice older than 18 months report an age-related decrease in wake and an increase in NREM across a 24-hour period^46–49^ while a few describe an increase in wakefulness and a decrease in NREM^50,51^. Another common finding, which is consistent with the human condition, is the increase in sleep fragmentation with age. Here, we report minimum effects of age on total sleep time as well as NREM sleep. However, we did observe an increase in fragmented sleep in our female cohort.

Results are also mixed regarding age-related changes in REM sleep, with some studies reporting no change with age^46,52^ or a decrease with aging^48,49^. While we only observed this decrease in REM in our male cohorts, an important caveat to consider is previous studies are almost exclusively conducted using male mice despite known biological sex differences in the sleep of C57Bl/6 mice^47,48,51,53^. We found an aging study utilizing mainly (but not exclusively) female mice, which revealed increased sleep fragmentation, shorter NREM bouts and no changes in REM with aging, which is consistent with our findings in our aged female cohort^52^.

It is also possible that the 18 month time point used here was too young to detect robust age-related changes in this strain of mice. Research by Hasan et al. 2012 observed that from 3 to 12 months, male C57Bl/6 mice exhibited a gradual decrease in time awake, with an increase in both NREM and REM sleep. However, by 24 months, mice showed a significant reversal of these with an increase in wake time and corresponding decrease in NREM and REM sleep to values surpassing those observed at 3 months^51^. We can speculate that our 18 month value would be somewhere between their 12 and 24 month values. This hypothetical value would likely correspond with their 3- or 6-month time point and could explain why we did not observe age-related significant changes. Age likely does influence sleep architecture in the C57Bl/6 mouse, but its effects could show parabolic changes as opposed to linear.

### *Centella Asiatica* improves sleep quantity and sleep maintenance

In this study, CAW was able to increase REM sleep duration in aged males, suggesting it can improve sleep quantity in older males but not females, further revealing sexual dimorphism in response to CAW. The observed REM increase was most apparent during the light portion of the light cycle. Because these animals are nocturnal, their sleep cycle typically shifts towards increased wakefulness and activity during the dark period so improved REM during the light portion is most ideal.

Increasing REM is unique as most medications approved for sleep disorders increase NREM sleep with minimal to no effect on REM sleep^54^. REM sleep is a unique physiological process associated with dreaming and is believed to maintain neuronal integrity, health, and function ^55^. While the exact function of REM sleep is heavily debated, it is proposed to serve as a house-keeping function of the brain, such as the removal of ROS^56^. There is strong evidence that impacted REM plays a prominent role in psychiatric diseases and, in cases of REM behavior sleep disorder, can be a significant risk factor and/or a precursor of neurodegenerative disorders^57–59^. A study showed that for every 1% decrease in REM sleep (but not NREM) there was an associated 9% increase in the risk of dementia^60^. Furthermore, decreases in REM sleep is associated with a greater risk of mortality as shown in 2 independent human cohorts with a 13% higher mortality rate for every 5% reduction in REM sleep^61^. Therefore, CAW increasing REM in our older male cohorts shows promise in slowing the effects of aging and improving/ maintaining brain health and function.

In addition to improving sleep architecture in older male cohorts, we also observed CAW resulting in a decrease in sleep fragmentation in our older females but not males. Sleep fragmentation alone has been shown to induce a myriad of negative health effects. Mice exposed to acute sleep fragmentation exhibit increased proinflammatory gene expressions in brain and peripheral organs^62,63^. Furthermore, rodent models of sleep fragmentation have been shown to accelerated Alzheimer disease (AD) progression in AD mice, with disease pathology occurring in several brain regions associated with sleep regulation and cognitive processing^64^. Furthermore, C57Bl/6 mice showed changes in aquaporin 4 density, which is indicative of glymphatic system impairment, following the same sleep fragmentation protocol^64^. The glymphatic system is a network of perivascular pathways that is known to support the clearance of brain interstitial proteins and is preferentially active during sleep. When this AQP4 channel is knocked out, fluid movement is lost and clearance of proinflammatory mediators therefore prolonging inflammation. By increasing ROS, sleep is further negatively impacted, and a positive feedback loop is perpetuating the problem.

### *Centella Asiatica* does not improve cognitive performance after sleep deprivation

Previously, it has been shown that CAW administration has improved cognitive performance in aged animals^17,31^. We wanted to evaluate if this effect was still present following sleep disruption therefore animals underwent 6 hours of sleep deprivation and were then tested 24 hours following the last training session for object recognition. Age-related impairments in object recognition /novel object discrimination have been observed in both rodents and humans^65–68^ however we failed to observe differences between our aged and young controls. Furthermore, the lack of improvements observed in NORT are curious as previous research using 20 month old CB6F1 in a similar paradigm revealed that both aged females and males receiving CAW spent more time with the novel object at 2 hours and 24 hours following the training phase^31^. Our lack of findings could be a result the added element of sleep deprivation.

The dual process hypothesis suggests that hippocampal dependent, declarative memory profits from SWS, whereas the consolidation of nondeclarative memory is supported by REM sleep^7–9^. In mice, memory consolidation primarily occurs within a few hours after learning, with the most critical period happening shortly after the initial encoding of information, often during periods of rest or sleep. Another study evaluating sleep deprivation was able to show improvements with NRF2 activating substances, however the sleep deprivation stage was prior to the initial habituating stage and mice were able to sleep directly after the first stage of training^69^. By preventing consolidation through sleep deprivation directly after training, we potentially hindered all groups ability to discriminate beyond what age alone could impair and beyond what CAW treatment could have helped. Furthermore, the amelioration of age-related cognitive decline by *Centella asiatica* extract varies by mode of administration. Recent work has shown significant improvements in cognitive function of aged mice following CAW administration through water while CAW administered through diet had limited behavioral effects potentially due to its low bioavailability^17,28,31,70,71^. Therefore, future studies should evaluate spatial memory following sleep deprivation with CAW administered through water prior to drawing conclusions.

### CAW may impact the antioxidant NRF2 pathway

Chronic insufficient sleep and sleep fragmentation are thought to aggravate the inflammatory response of nerve cells, which can contribute to existing neuroinflammation and increases in ROS, both conditions significantly associated with the development of neurodegenerative diseases^72–74^. ROS also increase during extended wakefulness or sleep deprivation, suggesting that sleep, in particular REM sleep, may function in ROS clearance^1,75^. Furthermore, with aging, ROS and sleep disruptions are known to both steadily increase^76,77^.

Hormones and different metabolic signaling pathways such as nuclear factor-kappa B (NF-kB) signaling, NRF2, silent information regulator 2-related protein (Sirt1) pathway and growth hormone axis are well established regulators of oxidative stress and inflammation in body implying they have influence on aging and longevity in mammals ^78–80^. In particular, the NRF2 pathway, is believed to regulate circadian rhythms and studies have shown that activation of this pathway can alleviate cognitive deficits incurred as a result of sleep deprivation ^81^. With increase in age, NRF2 signaling shows significant decline in efficiency^80,82^. Proper NRF2 pathway activity is suggested to be critical for maintaining healthy sleep quality including adequate REM sleep duration and stability^83^. Targeting these pathways present an opportunity to slow the progression of aging and improve the correlating symptoms including sleep impairments.

Therefore, NRF2 activation has become an attractive topic of research in sleep disturbances and disorders. CAW, a known NRF2 activator, has been shown to work on antioxidant gene expression in the cortex, hippocampus and cerebellum of aged mice^17^. CAW has also been shown to attenuate oxidative damage and decrease anxiety in sleep-deprived mice^84^ while other forms of NRF2 activation has been shown to alleviate cognitive impairments induced by sleep deprivation or sleep fragmentation^69,85–87^.

NRF2 activation, via CAW, and inactivation have shown differing effects in males compared to females in rodent studies of several different disease models^28,88–90^. Similarly, in a small human clinical study, *Centella asiatica* had a differential, but beneficial, cognitive effect between healthy middle-aged female and male participants. Several memory tasks showed improvements amongst males and females however, C. asiatica improved visual spatial thinking throughout the course of the trials in males only, while only females showed improvements in working memory, short term retrieval, and executive processing.^91^

While we also observed dimorphic effects of CAW, this differential influence based on sex could be a result of it impacting the only deficits to treat. In females, we did not see a change in REM with aging and with males we did not see more fragmented sleep with aging. Therefore, CAW can have a ceiling effect where the maximum REM sleep and lowest sleep fragmentation in females and males respectively is already being achieved therefore no improvements could be detected with CAW treatment. To better understand the dimorphic influence CAW has, future studies should investigate how CAW is metabolized in females versus males.

### Conclusions

Our results revealed that with age, quantity of sleep in males and quality of sleep in females declines. Daily treatment with CAW shows promise in improving these age-related sleep deficits. To gain a better understanding of CAWs influence on sleep quality in aging, future studies should utilize an older cohort and include the evaluation of sleep delta power which is often associated with sleep duration and intensity^92^.

## Acknowledgements

The data in this work was supported with resources and the use of facilities at the VA Portland Health Care System, the Portland VA Research Foundation, NIH U19 AT010829 to AS, NIH-NCCIH T32 AT002688 to LD, VA BLRD Merit Award 1I01BX006155 and DoD PRP HT9425-24-1-0775 to MML. The contents do not represent the views of the US Department of Veterans Affairs or the United States Government.

## Disclosure Statement

### Financial Disclosure

none.

### Non-financial Disclosure

none

## Notes

### Competing Interest Statement

The authors have declared no competing interest.

## Reference List

1. Maquet P, Peigneux P, Laureys S, Smith C. Be caught napping: you’re doing more than resting your eyes. Nat Neurosci. 2002;5(7):618–619. doi:10.1038/nn0702-618

2. Yoo SS, Hu PT, Gujar N, Jolesz FA, Walker MP. A deficit in the ability to form new human memories without sleep. Nat Neurosci. 2007;10(3):385–392. doi:10.1038/nn1851

3. Xie L, Kang H, Xu Q, et al. Sleep drives metabolite clearance from the adult brain. Science (1979). 2013;342(6156):373–377. doi:10.1126/science.1241224

4. Luyster FS, Strollo PJ, Zee PC, Walsh JK. Sleep: A health imperative. Sleep. 2012;35(6):727-734. doi:10.5665/sleep.1846

5. Ohayon MM, Carskadon MA, Guilleminault C, Vitiello M V. Meta-analysis of quantitative sleep parameters from childhood to old age in healthy individuals: Developing normative sleep values across the human lifespan. Sleep. 2004;27(7):1255–1273. doi:10.1093/sleep/27.7.1255

6. Walker MP, Stickgold R. Sleep, memory, and plasticity. Annu Rev Psychol. 2006;57:139-166. doi:10.1146/annurev.psych.56.091103.070307

7. Gais S, Born J. Declarative memory consolidation: Mechanisms acting during human sleep. Learning and Memory. 2004;11(6):679–685. doi:10.1101/lm.80504

8. Maquet P. The role of sleep in learning and memory. Science (1979). 2001;294(5544):1048–1052. doi:10.1126/science.1062856

9. Smith C. Sleep states and memory processes in humans: Procedural versus declarative memory systems. Sleep Med Rev. 2001;5(6):491–506. doi:10.1053/smrv.2001.0164

10. Drummond SPA, Brown GG. The effects of total sleep deprivation on cerebral responses to cognitive performance. Neuropsychopharmacology. 2001;25(5):S68–S73. doi:10.1016/S0893-133X(01)00325-6

11. Stricker JL, Brown GG, Wetherell LA, Drummond SPA. The impact of sleep deprivation and task difficulty on networks of fMRI brain response. Journal of the International Neuropsychological Society. 2006;12(5):591–597. doi:10.1017/S1355617706060851

12. Wetzel W, Wagner T, Balschun D. REM sleep enhancement induced by different procedures improves memory retention in rats. European Journal of Neuroscience. 2003;18(9):2611–2617. doi:10.1046/j.1460-9568.2003.02890.x

13. Harman D. Alzheimer’s disease pathogenesis: Role of aging. In: Annals of the New York Academy of Sciences. Vol 1067. Blackwell Publishing Inc.; 2006:454–460. doi:10.1196/annals.1354.065

14. Hayes MT. Parkinson’s Disease and Parkinsonism. American Journal of Medicine. 2019;132(7):802–807. doi:10.1016/j.amjmed.2019.03.001

15. Ureshino RP, Ramírez AL. Linking aging and animal models to neurodegeneration: The striatum, substantia nigra, and Parkinson’s disease. In: Assessments, Treatments and Modeling in Aging and Neurological Disease: The Neuroscience of Aging. Elsevier; 2021:539–552. doi:10.1016/B978-0-12-818000-6.00048-2

16. K.J G, J.A P, A.K G. Pharmacological Review on Centella asiatica: A Potential Herbal Cure-all. Indian J Pharm Sci. 2010;(October):546–556.

17. Gray NE, Harris CJ, Quinn JF, Soumyanath A. Centella asiatica modulates antioxidant and mitochondrial pathways and improves cognitive function in mice. J Ethnopharmacol. 2016;180:78–86. doi:10.1016/j.jep.2016.01.013

18. Wong JH, Barron AM, Abdullah JM. Mitoprotective Effects of Centella asiatica (L.) Urb.: Anti-Inflammatory and Neuroprotective Opportunities in Neurodegenerative Disease. Front Pharmacol. 2021;12(June):1–9. doi:10.3389/fphar.2021.687935

19. Haleagrahara N, Ponnusamy K. Neuroprotective effect of Centella asiatica extract (CAE) on experimentally induced parkinsonism in aged Sprague-Dawley rats. Journal of Toxicological Sciences. 2010;35(1):41–47. doi:10.2131/jts.35.41

20. Matthews DG, Caruso M, Murchison CF, et al. Centella asiatica improves memory and promotes antioxidative signaling in 5XFAD mice. Antioxidants. 2019;8(12). doi:10.3390/antiox8120630

21. Soumyanath A, Zhong YP, Henson E, et al. Centella asiatica extract improves behavioral deficits in a mouse model of Alzheimer’s disease: Investigation of a possible mechanism of action. Int J Alzheimers Dis. 2012;2012. doi:10.1155/2012/381974

22. Tiwari S, Singh S, Patwardhan K, Gehlot S, Gambhir IS. Effect of Centella asiatica on mild cognitive impairment (MCI) and other common age-related clinical problems Ayurveda Literature and Conceptual Insights View project Workshop on Scientific Writing for Ayurveda Scholars View project EFFECT OF CENTELLA ASI. Article in Digest Journal of Nanomaterials and Biostructures. 2008;3(4):215–220. https://www.researchgate.net/publication/201910063

23. Brinkhaus B, Lindner M, Schuppan D, Hahn EG. Chemical, pharmacological and clinical profile of the East Asian medical plant Centella asiatica. Phytomedicine. 2000;7(5):427–448. doi:10.1016/S0944-7113(00)80065-3

24. Bylka W, Znajdek-Awizeń P, Studzińska-Sroka E, Brzezińska M. Centella asiatica in cosmetology. Postepy Dermatol Alergol. 2013;30(1):46–49. doi:10.5114/pdia.2013.33378

25. Arpaia MR, Ferrone R, Amitrano M, Nappo C, Leonardo G, del Guercio R. Effects of Centella asiatica extract on mucopolysaccharide metabolism in subjects with varicose veins. Int J Clin Pharmacol Res. 1990;10(4):229–233. http://www.ncbi.nlm.nih.gov/pubmed/2150405

26. Bandopadhyay S, Mandal S, Ghorai M, et al. Therapeutic properties and pharmacological activities of asiaticoside and madecassoside: A review. J Cell Mol Med. 2023;27(5):593–608. doi:10.1111/jcmm.17635

27. Yang L, Marney L, Magana AA, et al. Quantification of caffeoylquinic acids and triterpenes as targeted bioactive compounds of Centella Asiatica in extracts and formulations by liquid chromatography mass spectrometry. Journal of Chromatography Open. 2023;4. doi:10.1016/j.jcoa.2023.100091

28. Gray NE, Hack W, Brandes MS, et al. Amelioration of age-related cognitive decline and anxiety in mice by Centella asiatica extract varies by sex, dose and mode of administration. Frontiers in Aging. 2024;5(May):1–14. doi:10.3389/fragi.2024.1357922

29. Lim MM, Elkind J, Xiong G, et al. Dietary Therapy Mitigates Persistent Wake Deficits Caused by Mild Traumatic Brain Injury. Sci Transl Med. 2013;5(215). doi:10.1126/scitranslmed.3007092

30. Jones CE, Opel RA, Kaiser ME, et al. Early-Life Sleep Disruption Increases Parvalbumin in Primary Somatosensory Cortex and Impairs Social Bonding in Prairie Voles. Vol 5.; 2019. https://www.science.org

31. Gray NE, Zweig JA, Caruso M, et al. Centella asiatica increases hippocampal synaptic density and improves memory and executive function in aged mice. Brain Behav. 2018;8(7):1–11. doi:10.1002/brb3.1024

32. Dijk DJ, Duffy JF, Czeisler CA. CONTRIBUTION OF CIRCADIAN PHYSIOLOGY AND SLEEP HOMEOSTASIS TO AGE-RELATED CHANGES IN HUMAN SLEEP. Chronobiol Int. 2000;17(3):285–311. doi:10.1081/CBI-100101049

33. Vitiello M V. Sleep in Normal Aging. Sleep Med Clin. 2006;1(2):171–176. doi:10.1016/j.jsmc.2006.04.007

34. Brassington GS, King AC, Bliwise DL. Sleep problems as a risk factor for falls in a sample of community-dwelling adults aged 64-99 years. J Am Geriatr Soc. 2000;48(10):1234–1240. doi:10.1111/j.1532-5415.2000.tb02596.x

35. Kaufmann CN, Canham SL, Mojtabai R, et al. Insomnia and health services utilization in middle-Aged and older adults: Results from the health and retirement study. Journals of Gerontology - Series A Biological Sciences and Medical Sciences. 2013;68(12 A):1512-1517. doi:10.1093/gerona/glt050

36. Newman AB, Enright PL, Manolio TA, Haponik EF, Wahl PW. Sleep Disturbance, Psychosocial Correlates, and Cardiovascular Disease in 5201 Older Adults: The Cardiovascular Health Study. J Am Geriatr Soc. 1997;45(1):1-7. doi:10.1111/j.1532-5415.1997.tb00970.x

37. Cappuccio FP, Cooper D, Delia L, Strazzullo P, Miller MA. Sleep duration predicts cardiovascular outcomes: A systematic review and meta-analysis of prospective studies. Eur Heart J. 2011;32(12):1484–1492. doi:10.1093/eurheartj/ehr007

38. Cappuccio FP, Taggart FM, Kandala NB, et al. Meta-analysis of short sleep duration and obesity in children and adults. Sleep. 2008;31(5):619–626. doi:10.1093/sleep/31.5.619

39. Dzierzewski JM, Dautovich N, Ravyts S. Sleep and Cognition in Older Adults. Sleep Med Clin. 2018;13(1):93–106. doi:10.1016/j.jsmc.2017.09.009

40. Dzierzewski JM, Dautovich ND. Who Cares about Sleep in Older Adults? Clin Gerontol. 2018;41(2):109–112. doi:10.1080/07317115.2017.1421870

41. Knutson KL, Van Cauter E. Associations between Sleep Loss and Increased Risk of Obesity and Diabetes. Ann N Y Acad Sci. 2008;1129(1):287–304. doi:10.1196/annals.1417.033

42. Lim J, Dinges DF. A meta-analysis of the impact of short-term sleep deprivation on cognitive variables. Psychol Bull. 2010;136(3):375–389. doi:10.1037/a0018883

43. Webb WB, Levy CM. Age, Sleep Deprivation, and Performance. Psychophysiology. 1982;19(3):272-276. doi:10.1111/j.1469-8986.1982.tb02561.x

44. Scott AJ, Webb TL, Martyn-St James M, Rowse G, Weich S. Improving sleep quality leads to better mental health: A meta-analysis of randomised controlled trials. Sleep Med Rev. 2021;60:101556. doi:10.1016/j.smrv.2021.101556

45. Haack M, Serrador J, Cohen D, Simpson N, Meier-Ewert H, Mullington JM. Increasing sleep duration to lower beat-to-beat blood pressure: A pilot study. J Sleep Res. 2013;22(3):295–304. doi:10.1111/jsr.12011

46. Soltani S, Chauvette S, Bukhtiyarova O, et al. Sleep–Wake Cycle in Young and Older Mice. Front Syst Neurosci. 2019;13. doi:10.3389/fnsys.2019.00051

47. Panagiotou M, Vyazovskiy V V., Meijer JH, Deboer T. Differences in electroencephalographic non-rapid-eye movement sleep slow-wave characteristics between young and old mice. Sci Rep. 2017;7(January):1–12. doi:10.1038/srep43656

48. Masuda K, Katsuda Y, Niwa Y, Sakurai T, Hirano A. Analysis of circadian rhythm components in EEG/EMG data of aged mice. Front Neurosci. 2023;17(May):1–11. doi:10.3389/fnins.2023.1173537

49. Wimmer ME, Rising J, Galante RJ, Wyner A, Pack AI, Abel T. Aging in mice reduces the ability to sustain sleep/wake states. PLoS One. 2013;8(12):4–12. doi:10.1371/journal.pone.0081880

50. Eleftheriou BE, Zolovick AJ, Elias MF. Electroencephalographic Changes with Age in Male Mice. Gerontology. 1975;21(1):21–30. doi:10.1159/000212027

51. Hasan S, Dauvilliers Y, Mongrain V, Franken P, Tafti M. Age-related changes in sleep in inbred mice are genotype dependent. Neurobiol Aging. 2012;33(1):195.e13–195.e26. doi:10.1016/j.neurobiolaging.2010.05.010

52. Tsuji S, Brace CS, Yao R, et al. Sleep–wake patterns are altered with age, Prdm13 signaling in the DMH, and diet restriction in mice. Life Sci Alliance. 2023;6(6). doi:10.26508/lsa.202301992

53. Mannino GS, Green TRF, Murphy SM, Donohue KD, Opp MR, Rowe RK. The importance of including both sexes in preclinical sleep studies and analyses. Sci Rep. 2024;14(1):23622. doi:10.1038/s41598-024-70996-1

54. Atkin T, Comai S, Gobbi G. Drugs for insomnia beyond benzodiazepines: Pharmacology, clinical applications, and discovery. Pharmacol Rev. 2018;70(2):197–245. doi:10.1124/pr.117.014381

55. Nir Y, Tononi G. Dreaming and the brain: from phenomenology to neurophysiology. Trends Cogn Sci. 2010;14(2):88–100. doi:10.1016/j.tics.2009.12.001

56. Mallick BN, Singh A. REM sleep loss increases brain excitability: Role of noradrenalin and its mechanism of action. Sleep Med Rev. 2011;15(3):165–178. doi:10.1016/j.smrv.2010.11.001

57. Shen Y, Yu WB, Shen B, et al. Propagated a-synucleinopathy recapitulates REM sleep behaviour disorder followed by parkinsonian phenotypes in mice. Brain. 2020;143(11):3374–3392. doi:10.1093/BRAIN/AWAA283

58. Marion MH, Qurashi M, Marshall G, Foster O. Is REM sleep Behaviour Disorder (RBD) a risk factor of dementia in idiopathic Parkinson’s disease? J Neurol. 2008;255(2):192–196. doi:10.1007/s00415-008-0629-9

59. Galbiati A, Verga L, Giora E, Zucconi M, Ferini-Strambi L. The risk of neurodegeneration in REM sleep behavior disorder: A systematic review and meta-analysis of longitudinal studies. Sleep Med Rev. 2019;43:37–46. doi:10.1016/j.smrv.2018.09.008

60. Pase MP, Himali JJ, Grima NA, et al. Sleep Architecture and the Risk of Incident Dementia in the Community.; 2017.

61. Leary EB, Watson KT, Ancoli-Israel S, et al. Association of Rapid Eye Movement Sleep with Mortality in Middle-aged and Older Adults. JAMA Neurol. 2020;77(10):1241–1251. doi:10.1001/jamaneurol.2020.2108

62. Mishra I, Pullum KB, Thayer DC, et al. Chemical sympathectomy reduces peripheral inflammatory responses to acute and chronic sleep fragmentation. Am J Physiol Regul Integr Comp Physiol. 2020;318:781–789. doi:10.1152/ajpregu.00358.2019.-Sleep

63. Dumaine JE, Ashley NT. Acute sleep fragmentation does not alter pro-inflammatory cytokine gene expression in brain or peripheral tissues of leptin-deficient mice. PeerJ. 2018;2018(2). doi:10.7717/peerj.4423

64. Vasciaveo V, Iadarola A, Casile A, et al. Sleep fragmentation affects glymphatic system through the different expression of AQP4 in wild type and 5xFAD mouse models. Acta Neuropathol Commun. 2023;11(1). doi:10.1186/s40478-022-01498-2

65. Marshall HJ, Pezze MA, Fone KCF, Cassaday HJ. Age-related differences in appetitive trace conditioning and novel object recognition procedures. Neurobiol Learn Mem. 2019;164. doi:10.1016/j.nlm.2019.107041

66. Kaviani E, Rahmani M, Kaeidi A, et al. Protective effect of atorvastatin on D-galactose-induced aging model in mice. Behavioural Brain Research. 2017;334(April):55–60. doi:10.1016/j.bbr.2017.07.029

67. Merriman NA, Ondřej J, Roudaia E, O’Sullivan C, Newell FN. Familiar environments enhance object and spatial memory in both younger and older adults. Exp Brain Res. 2016;234(6):1555–1574. doi:10.1007/s00221-016-4557-0

68. Singh P, Thakur MK. Reduced recognition memory is correlated with decrease in DNA methyltransferase1 and increase in histone deacetylase2 protein expression in old male mice. Biogerontology. 2014;15(4):339–346. doi:10.1007/s10522-014-9504-5

69. Wang W, Yang L, Liu T, Wang J, Wen A, Ding Y. Ellagic acid protects mice against sleep deprivation-induced memory impairment and anxiety by inhibiting TLR4 and activating Nrf2. Aging. 2020;12(11):10457–10472. doi:10.18632/aging.103270

70. Zweig JA, Brandes MS, Brumbach BH, et al. Loss of NRF2 accelerates cognitive decline, exacerbates mitochondrial dysfunction, and is required for the cognitive enhancing effects of Centella asiatica during aging. Neurobiol Aging. 2021;100(5):48–58. doi:10.1016/j.neurobiolaging.2020.11.019

71. Speers AB, Wright KM, Brandes MS, et al. Mode of administration influences plasma levels of active Centella asiatica compounds in 5xFAD mice while markers of neuroinflammation remain unaltered. Front Neurosci. 2024;18. doi:10.3389/fnins.2024.1277626

72. McAlpine CS, Kiss MG, Zuraikat FM, et al. Sleep exerts lasting effects on hematopoietic stem cell function and diversity. J Exp Med. 2022;219(11). doi:10.1084/jem.20220081

73. Singh A, Kukreti R, Saso L, Kukreti S. Oxidative stress: A key modulator in neurodegenerative diseases. Molecules. 2019;24(8). doi:10.3390/molecules24081583

74. Zhang W, Xiao D, Mao Q, Xia H. Role of neuroinflammation in neurodegeneration development. Signal Transduct Target Ther. 2023;8(1). doi:10.1038/s41392-023-01486-5

75. Reimund E. The Free Radical Flux Theory of Sleep. Vol 43.; 1994.

76. Maldonado E, Morales-Pison S, Urbina F, Solari A. Aging Hallmarks and the Role of Oxidative Stress. Antioxidants. 2023;12(3). doi:10.3390/antiox12030651

77. Bliwise DL. Sleep in Normal Aging and Dementia. Sleep. 1993;16(1):40–81. doi:10.1093/sleep/16.1.40

78. Bartke A. Pleiotropic effects of growth hormone signaling in aging. Trends in Endocrinology & Metabolism. 2011;22(11):437–442. doi:10.1016/j.tem.2011.07.004

79. García-García VA, Alameda JP, Page A, Casanova ML. Role of nf-κb in ageing and age-related diseases: Lessons from genetically modified mouse models. Cells. 2021;10(8):1–29. doi:10.3390/cells10081906

80. Chen C, Zhou M, Ge Y, Wang X. SIRT1 and aging related signaling pathways. Mech Ageing Dev. 2020;187(January):111215. doi:10.1016/j.mad.2020.111215

81. Wible RS, Ramanathan C, Sutter CH, et al. NRF2 regulates core and stabilizing circadian clock loops, coupling redox and timekeeping in mus musculus. Elife. 2018;7:1–27. doi:10.7554/eLife.31656

82. Zhang H, Davies KJA, Forman HJ. Oxidative stress response and Nrf2 signaling in aging. Free Radic Biol Med. 2015;88(Part B):314–336. doi:10.1016/j.freeradbiomed.2015.05.036

83. Davinelli S, Medoro A, Savino R, Scapagnini G. Sleep and Oxidative Stress: Current Perspectives on the Role of NRF2. Cell Mol Neurobiol. 2024;44(1). doi:10.1007/s10571-024-01487-0

84. Chanana P, Kumar A. Possible Involvement of Nitric Oxide Modulatory Mechanisms in the Neuroprotective Effect of Centella asiatica Against Sleep Deprivation Induced Anxiety Like Behaviour, Oxidative Damage and Neuroinflammation. Phytotherapy Research. 2016;30(4):671–680. doi:10.1002/ptr.5582

85. Wang W, Yang L, Liu T, et al. Corilagin ameliorates sleep deprivation-induced memory impairments by inhibiting NOX2 and activating Nrf2. Brain Res Bull. 2020;160:141–149. doi:10.1016/j.brainresbull.2020.03.010

86. Li Y, Xie Z, Luo X, et al. Farnesol Exerts Protective Effects against Chronic Sleep Deprivation-Induced Cognitive Impairment via Activation SIRT1/Nrf2 Pathway in the Hippocampi of Adult Mice. Mol Nutr Food Res. 2023;67(11). doi:10.1002/mnfr.202200735

87. Qiu X, Li L, Wei J, et al. The protective role of Nrf2 on cognitive impairment in chronic intermittent hypoxia and sleep fragmentation mice. Int Immunopharmacol. 2023;116. doi:10.1016/j.intimp.2023.109813

88. Kerns ML, Hakim JMC, Zieman A, Lu RG, Coulombe PA. Sexual Dimorphism in Response to an NRF2 Inducer in a Model for Pachyonychia Congenita. Journal of Investigative Dermatology. 2018;138(5):1094–1100. doi:10.1016/j.jid.2017.09.054

89. Yin Y, Corry KA, Loughran JP, Li J. Moderate Nrf2 Activation by Genetic Disruption of Keap1 Has Sex-Specific Effects on Bone Mass in Mice. Sci Rep. 2020;10(1). doi:10.1038/s41598-019-57185-1

90. McCallum RT, Thériault RK, Manduca JD, et al. Nrf2 activation rescues stress-induced depression-like behaviour and inflammatory responses in male but not female rats. Biol Sex Differ. 2024;15(1). doi:10.1186/s13293-024-00589-0

91. Dev RDO, Mohamed S, Hambali Z, Samah BA. Comparison on cognitive effects of Centella asiatica in healthy middle age female and male volunteers. European Journal of Scientific Research. 2009;31(4):553–565.

92. Long S, Ding R, Wang J, Yu Y, Lu J, Yao D. Sleep Quality and Electroencephalogram Delta Power. Front Neurosci. 2021;15. doi:10.3389/fnins.2021.803507

